# Positive supercoiling favors transcription elongation through lac repressor-mediated DNA loops

**DOI:** 10.1101/2021.03.02.433568

**Authors:** Wenxuan Xu, Yan Yan, Irina Artsimovitch, David Dunlap, Laura Finzi

**Affiliations:** Physics Department, Emory University, Atlanta, GA, USA; Department of Microbiology, Ohio State University, Columbus, OH, USA

## Abstract

Some proteins, like the *lac* repressor (LacI), mediate long-range loops that alter DNA topology and create torsional barriers. During transcription, RNA polymerase generates supercoiling that may facilitate passage through such barriers. We monitored *E. coli* RNA polymerase progress along templates in conditions that prevented, or favored, LacI-mediated DNA looping. Tethered particle motion measurements revealed that RNA polymerase paused longer at unlooped LacI obstacles or those barring entry to a loop than those barring exit from the loop. Enhanced dissociation of a LacI roadblock by the positive supercoiling generated ahead of a transcribing RNA polymerase within a torsion-constrained DNA loop may be responsible. In support of this, RNA polymerase transcribed a looped segment more slowly than an unlooped segment and paused more briefly at LacI obstacles on positively supercoiled templates, especially under increased tension (torque). Positive supercoiling propagating ahead of polymerase appears to facilitate elongation along topologically-complex, protein-coated templates.

## Introduction

DNA in the cell is complexed with proteins and adopts a highly compact structure including protein-mediated loops and supercoiling (1-11). Genome-bound proteins can be roadblocks that hinder elongation by RNA polymerase (RNAP) during transcription (12). The strength of such roadblocks may vary if the protein in question wraps DNA or secures a DNA loop. Several recent investigations have been conducted to understand how RNA polymerase navigates through nucleosomes (13-15). Nucleosomes contain histones which interact with DNA non-specifically and are substrates for combinatorial post-translational modifications that regulate chromatin remodelling and transcription of DNA. However, many transcription factors from organisms spanning all kingdoms shape genomes and influence transcription without such extensive chemical modification. Often, their activity is regulated by concentration, DNA supercoiling, and for the many which recognize specific sites on DNA, by the presence of multiple binding sites with different affinities. These transcription factors may bend, wrap, bridge and loop DNA segments (5, 16-24). The effects of these topologies have not been addressed in earlier studies on transcription roadblocks *in vivo* and are just beginning to be investigated *in vitro (25)*.

In this study, the *Escherichia coli* lac repressor (LacI) was used as a transcriptional roadblock (26). The bivalent LacI tetramer binds specific sites (operators) with up to nanomolar affinity, depending on the sequence, and can either bind to one high-affinity operator or to two operators simultaneously, mediating a DNA loop. The strength of the roadblock depends on the affinity for the binding site(s), on tethering, which increases the effective local concentration of the protein near a binding site(s) (27), and on the loop thermodynamic stability. Using a DNA template containing a weak binding site (O2) near a promoter and a high-affinity binding site (O1) further downstream, it was shown that LacI bound to O2 was a much stronger roadblock when securing a loop between the two operators (25). However, transcription of the loop segment by RNAPs that bypass the promoter-proximal LacI roadblock and the effect of the loop on the roadblocking capacity of promoter-distal roadblock have not been previously investigated. Thus, we used the tether particle motion (TPM) technique, to monitor the process of transcription through a LacI-mediated loop.

Surprisingly, we found that RNAP paused for long times within the loop region and that the loop weakened the promoter-distal roadblock. According to the twin-domain model (28), an elongating RNAP generates negative supercoiling upstream and positive supercoiling downstream. In addition, a protein-mediated loop is a barrier against diffusion of supercoiling (29). Therefore, we hypothesized that while negative supercoiling behind RNAP inside the loop stabilizes the binding of LacI at the promoter-proximal operator (30, 31), the positive supercoiling accumulated ahead of RNAP may destabilize LacI at the promoter-distal binding site, reducing its strength as a roadblock. Although it has been shown that transcription-generated positive supercoiling can destabilize nucleosomes (32), it is still not clear how supercoiling affects the binding affinity of other transcription factors or the topologies these factors induce. Thus, we used magnetic tweezers (MTs) to test the effect of supercoiling on LacI roadblocks. We found that RNAP paused more briefly in front of an O1-bound LacI when the DNA template was positively supercoiled than in the absence of supercoiling. This observation supports the idea that transcription-induced supercoiling within a LacI-mediated loop stabilizes LacI binding to the operator behind while destabilizing LacI binding to the operator ahead of the transcription complex.

In summary, our study reveals complex effects of protein-stabilized loops on the kinetics of RNA chain elongation. The LacI-mediated loops lengthen the pause time in front of the promoter-proximal binding site, shorten the pause time in front of the promoter-distal binding site, and increase the frequency and duration of pausing within the loop. We conclude that DNA loops can be potent transcription roadblocks that can temporarily sequester RNAP, until positive supercoiling build up breaks the loop and enables RNAP escape. The significance of these findings is that *in vivo*, should RNAP elongation be hindered in loops due to protein-mediated long-range interactions, the enzyme has an intrinsic ability to generate sufficient positive supercoiling to free itself from the loop and complete the synthesis of the nascent RNA chain.

## Materials and Methods

### Preparation of DNA constructs

DNA tethers for TPM experiments (Figure S1A) were amplified in PCR reactions with plasmid templates containing various combinations of LacI operators. Tethers with O1 and O2 in the proximal and distal position from the promoter, respectively, were amplified from template pWX_12_400 (see SI for sequence). A tether with O2 and O1 in the opposite order was amplified from pZV_21_400 (33). Constructs with a single O1 operator were amplified from pRS_1N_400 (see SI for sequence). The reactions contained dNTPs (Fermentas-Thermo Fisher Scientific Inc., Pittsburgh, PA, USA), biotin-labeled, or unlabeled anti-sense primers (Eurofins Genomics, Louisville, KY or Integrated DNA Technologies, Coralville, IA, USA) and Taq DNA Polymerase (New England BioLabs, Ipswich, MA, USA).

DNA tethers used in the MTs measurements were built by ligating a 4098 bp-long main fragment containing the promoter-proximal O1 operator to a ∼150 bp-long multiple biotin-labeled DNA fragment at the promoter-distal end with T7 DNA ligase (New England Bio-Labs, Ipswich, MA, USA). The main fragment was amplified from pRS_1N_400 using an equimolar dNTP mix and a primer containing an ApaI restriction site, digested with ApaI, and purified on a PCR clean up column. A 302 bp-long biotin-labeled amplicon was also produced using dATP, dCTP, dGTP, dTTP (Fermentas-Thermo Fisher Scientific Inc., Pittsburgh, PA, USA) and biotin-11-dUTP (Invitrogen, Life Technologies, Grand Island, NY, USA) in a molar ratio of 1:1:1:0.7:0.3. Digestion of this fragment with ApaI generated ∼150 bp biotin-labeled “tail” fragments for ligation. The biotinylated “tail” fragment anchored the tether, through multiple attachment sites, to a streptavidin-coated bead, allowing torque to be applied to the DNA with the magnetic tweezer (Figure S1B).

On all constructs, RNAP could be stalled 22 bp from the transcription start site (TSS) by withholding CTP.

Table S1 summarizes the name of the plasmids, primers, and restriction enzymes, as well as the sequence of the primers used.

### Preparation of RNAP

Doubly-HA tagged *E. coli* RNAP was used in both TPM and MTs experiments. This enzyme was purified as described previously (34).

### Microchamber preparation

The bottom coverslip of the microchamber (Fisherbrand, Thermo Fisher Scientific, Waltham, MA, USA) supported a parafilm gasket produced with a laser cutter (Universal Laser Systems, VLS 860, Middletown, CT) with a central observation area connected through narrow inlet and outlet channels to inlet and outlet reservoirs, extending beyond the edges of the top coverslip, (Figure S2). The coverslips with parafilm assembly were heated to seal its components together and form the microchamber. The narrow inlet and outlet reduced evaporation of buffer, while the triangular shape of the chamber, confined the reaction in a relatively small volume and provided a gradient of tether densities to optimize throughput.

The entire sample preparation was performed at room temperature (∼25 °C) and materials were kept on ice. Beads were first washed with phosphate buffered saline (PBS), and then washed and resuspended in transcription buffer (TXB: 20 mM Tris-glutamate (pH 7.4), 10 mM magnesium glutamate, 50 mM potassium-glutamate, 0.2 mg/mL α-casein (Sigma-Aldrich, St. Louis, MO), 1 mM DTT). The chambers were first incubated with reference beads resuspended in basic incubation buffer (BIB) (20 mM Tris-Cl (pH 7.4), 50 mM KCl, 1 mM DTT) for 5 minutes to let some beads adhere to the surface and serve as references during data acquisition and analysis. The chambers were then incubated with 10 μg/ml purified Anti-HA 11 Epitope tag antibody (BioLegend, San Diego, CA, USA) in BIB at room temperature for 1 hour. They were then passivated with BIB supplemented with 6 mg/ml α-casein at 4 °C overnight. 60 nM doubly-HA tagged *E. coli* RNA polymerase were then drawn into the chamber and incubated 30 min at room temperature to let the HA-labeled RNA polymerase bind to the anti-HA coated surface. 10 nM DNA template, 50 µM GpA (initiating dinucleotide, TriLink Bio Technologies, San Diego, CA, USA), and 10 µM ATP/UTP/GTP in TXB were introduced into the chamber to produce the stalled elongation complexes (SECs). For TPM experiments, the end of the DNA far from the promoter was labeled with beads (0.32 μm diameter, streptavidin-coated polystyrene beads (Roche Life Science, Indianapolis, IN, USA)) by flowing in the microchamber 0.03 mg/mL beads resuspended in TXB and letting them incubate for 10 min. For MTs experiments, instead, a 0.02 mg/ml solution of 1.0 μm diameter, streptavidin-coated superparamagnetic beads (Dynabead MyOne Streptavidin T1, Invitrogen, Grand Island, NY)) was incubated for 5 min. Both types of beads were washed in PBS twice and resuspended in TXB before introducing them into the chamber. The experimental construct for TPM/MTs is schematically illustrated in Figure S1. The extension of the DNA tether was monitored before, to control the original tether length, and after introducing 1 mM NTPs with/without LacI in TXB.

### The tethered particle motion technique and data analysis

The tethered particle motion (TPM) technique has been described in detail elsewhere (35). All TPM experiments were conducted at room temperature. The lamp of the microscope was turned on approx. 1-2 hours before recording the motion of the beads to thermally equilibrate the microscope. Five immobilized beads were used to calculate the drift, which was then subtracted from the tethered particle data (35, 36). The absolute *x, y* positions of each bead were recorded at 50 Hz with a customized Labview (National Instruments, Austin, TX, USA) software. The excursion of each tether was then calculated as 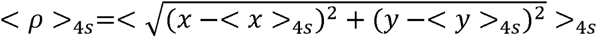, in which (< *x* >_4*s*_, < *y* >_4*s*_ i represents a four-second moving average of the coordinates that reveals the anchor point of the bead. The average excursion <ρ> was based on a four-second moving average of the variance. Changes in the bead excursion reflected changes in the DNA tether length (37-39). The recorded *x, y* data were then analyzed for symmetry and initial tether length prior to the start of transcription. Beads that exhibited (*x, y*) position distributions with a ratio of the major to minor axes greater than 1.07, were discarded, since they were likely to be tethered by multiple DNA molecules(35). The tether length was determined using a calibration curve relating tether length to <ρ^2^> values (Figure S3). Tethers with anomalous length were excluded.

### Magnetic Tweezers

Our magnetic tweezers parameters have been previously described (40, 41). All MTs experiments were conducted at room temperature using a protocol similar to what already described (33). Twisting the DNA tether, anchored via a stalled RNAP (Figure S1), produced plectonemes under low tension (<1 pN) after reaching the critical torque value. An extension versus twist curve displayed the extension (ΔZ) of DNA tether versus the number of magnet turns (Figure S4, left) and was used to distinguish beads tethered by intact, single dsDNA tethers from linked to nicked molecules or multiply DNA tethers. Prior to adding NTPs to initiate transcription, tethers were twisted by -19 turns and the extension was recorded for about 1 minute. Then 1 mM NTPs with or without 10 nm LacI in TXB were introduced to resume transcription.

### Estimation of RNAP progress within the loop

When a transcription elongation complex (TEC) is trapped inside a loop, torque will accumulate quickly, and a torque of +11 pN·nm ahead, or -11 pN·nm behind RNAP has been shown to stall its progress (42). Torque will quickly accumulate and stall TECs close to either of the LacI binding sites sealing the loop but less so when the enzyme is in the middle of the loop. At this location, the torque in the flanking DNA can be expressed as:

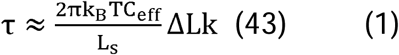

in which L_s_ is the contour length of the flanking DNA. C_eff_ is the effective twist persistence length (twist modulus/k_B_T) determined as 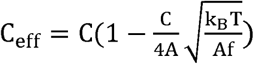 (43), where C=100nm, A=50nm, and the tension, f, in this case is 0.45 pN. ΔLk is the linking number change in the DNA. In this study 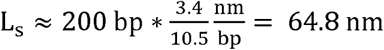, C_eff_ ≈ 63 nm, k_B_T≈ 4.1 pN·nm. Therefore, the maximum change in linking number that RNAP can induce before stalling (|τ| ≈ 11 pN·nm), is 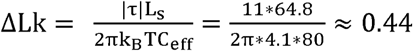 turns, which corresponds to 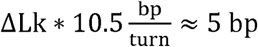. Thus, 5 bp is the furthest that RNAP should be expected to transcribe before stalling when it operates inside a ∼400 bp loop, and most likely occurs when RNAP is halfway between the two operators.

### Estimation of torque at the O1 pause site

Mosconi et al. (44) have shown that under 0.45 pN and 0.8 pN of tension C_eff_ ≈ 63 nm and 72 nm, respectively. Since the initial twist of the DNA tether may be arbitrarily set and RNAP (+) supercoils the downstream DNA, a DNA tether can be configured to have ΔLk = 5 just as a TEC reaches O1. The length of the DNA tether at that point is 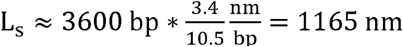, and equation (1) yields a torque of τ _0.45_ ≈ 7.0 pN·nm, τ _0.8_ ≈ 8.0 pN·nm at tensions of 0.45 or 8 pN, respectively. Instead, under 0.25 pN tension, ΔLk = 5 occurs beyond the buckling transition, a phase in which the torque in the plectonemic DNA remains constant at approximately 4 pN·nm (44).

## Results

### Monitoring elongation through LacI-mediated loops with tethered particle motion

First, TPM was used to study elongation through LacI-mediated loops. As RNAP transcribed the DNA template containing two binding sites for the LacI protein (Figure 1A), three scenarios were possible: RNAP might encounter an unencumbered binding site, a binding site bound by LacI in unlooped DNA, or a LacI bridging two operators to secure a DNA loop. In addition, during the progress of RNAP, LacI might randomly bind to/dissociate from either the promoter-proximal or - distal binding site, O_prox_ and O_dist_, respectively, producing/breaking intermittent loops. In data that satisfied screening criteria, Figure 1B for example, prior to addition of all four nucleotides the average excursion of the bead remained constant at a value consistent with the DNA tether length. Addition of all NTPs without and with LacI, indicated by the blue arrow in Figures 1B and C, caused a short-lived disturbance, which was deleted from the record. Then, RNAP resumed elongation producing a progressive decrease in tether length that continued uninterrupted to the end of the template, unless LacI was present. Control experiments with 1 mM NTPs revealed no pausing by RNAP at either of the two LacI binding sites in the absence of LacI (Figure 1B, left). In contrast, in the presence of 10 nm LacI, RNAP clearly paused in front of O_dist_ on a DNA template containing only this LacI binding site (Figure 1B, middle) or paused in front of both O_prox_ and O_dist_ on a DNA template containing two LacI binding sites (Figure 1B, right).

**Fig. 1.**
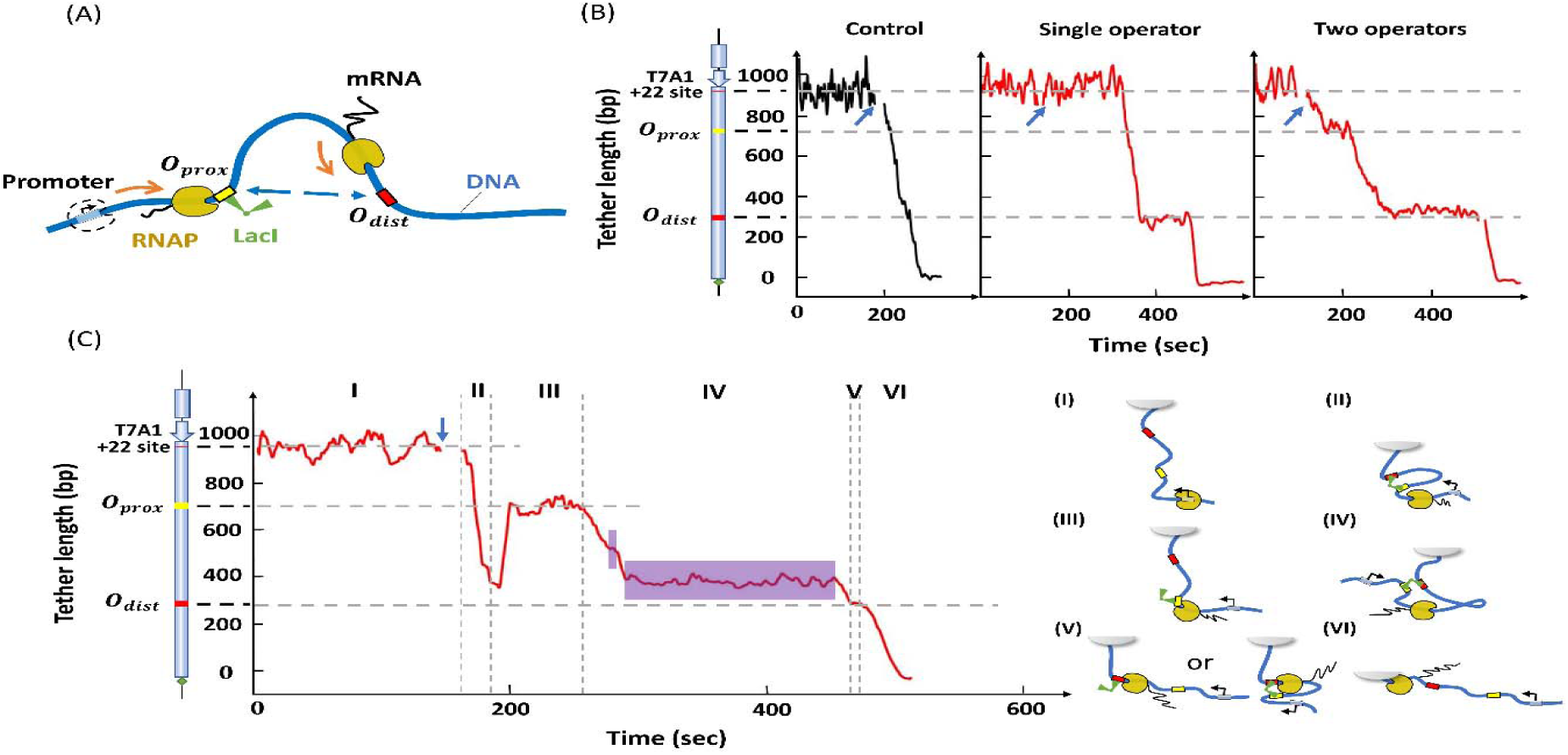
RNAP can transcribe through a LacI-mediated loop after a pause. (A) The transcription elongation complex is depicted at different points along the template. RNAP (yellow with black nascent RNA), DNA (thick blue line), LacI binding sites (yellow and red bars on blue DNA) and LacI (green, V-shape). Curved orange arrows indicate the direction of transcription starting at the promoter. A single LacI may bridge the two binding sites and induce the formation of a DNA loop. (B) Representative transcription records. Left: control record without LacI. Center: transcription in the presence of 10 nm LacI and only the distal operator. Right: transcription in the presence of 10 nm LacI and two LacI binding sites. At this concentration, each binding site is likely to be occupied by a different LacI tetramer; no looping is likely. The blue arrows indicate the time at which NTPs were introduced. (C) Transcription in the presence of 0.2 nm LacI, conditions that promote looping. The vertical dash lines identify six intervals (I - VI) in the progress of RNAP along the DNA template and the cartoons on the right depict the likely conformation of the transcription elongation complex in each interval. The purple areas in region IV indicate random pauses between operators. The cartoon on the left of the y-axis in panels B and C shows the features of the DNA template used in TPM measurements. From top to bottom: a T7A1 promoter, a stall site at +22, a promoter-proximal binding site (O_prox_) and promoter-distal binding site (O_dist_). The dashed horizontal lines indicate the position of the LacI binding site(s) in the construct, and the expected location of pauses in the record.

The probability of looping in a DNA template containing two binding sites can be adjusted by varying the LacI concentration (27). For a 400-bp loop between the O2 and O1 binding sites, a maximum looping probability of ∼40% could be achieved with 0.2 nM LacI, while 10 nM LacI decreased the probability to ∼6% as the two binding sites became occupied by different LacI tetramers (Figure S5). Indeed, with 10 nM LacI, RNAP paused at positions corresponding to the promoter-proximal and -distal LacI binding sites and no loops were observed (Figure 1B, right).

However, with 0.2 nM LacI, loops occurred during transcription (Figure 1C) and the pattern was more complex. The record can be divided into six intervals (I-IV). In interval (I) the DNA tether length is constant before introducing 1 mM NTPs and 0.2 nM LacI. Transcription begins shortly after introducing NTPs and in interval II, a loop forms, but it ruptures as RNAP approaches O_prox_; since RNAP pauses at this location, LacI likely remains bound to O_prox_. In interval (III), RNAP paused at O_prox_ for approximately 50 s. In interval (IV), RNAP proceeded to transcribe the segment between the LacI binding sites and paused at random locations (purple areas) before reaching the distal operator. Traces without loop formation (Figure 1B) did not reveal any sequence-induced pauses. Thus, we infer that pauses within the loop may be induced by the accumulation of supercoiling that takes place inside a loop, as depicted in Figure 1C cartoon IV (see also “Materials and Methods/Estimation of RNAP pauses within the loop”). Elongation resumed after loop breakdown and RNAP transcribed to O_dist_. In interval (V), RNAP paused at O_dist_ indicating that LacI was still associated with this operator, then continued transcription in interval (VI), finally reaching the end of the template. This record illustrates how transcription through a loop can be monitored using TPM.

### RNAP pauses longer at entry to than exit from LacI-loops

Previous work showed that LacI bound to a promoter-proximal O2 operator blocked transcription more effectively when securing a loop (33), a conclusion drawn from an analysis of the distance traveled by RNAP before immobilization for static AFM imaging. In those experiments, stalled transcription elongation complexes, reactivated upon the addition of the missing ribonucleotide triphosphate, transcribed segments as long as the entire 906 bp template in 60 seconds, and were even observed inside the Lac-mediated loop. To measure the time required for transcription of the loop and determine whether looping affected the passage of RNAP through bound LacI, the pause times of RNAP at O1 or O2 binding sites in the O_prox_ position on unlooped templates (Figure 1B, right) were compared with those in looped templates (Figure S6, interval III). On unlooped templates, pauses at proximal and distal O1 (O2) operators occupied by LacI were similar and their pause time were aggregated into figure 2. By contrast, formation of a loop had profound effects on RNAP elongation. RNAP paused longer at promoter-proximal LacI-O1 or LacI-O2 obstacles that were part of a loop (Figure 2A, left).

**Fig. 2.**
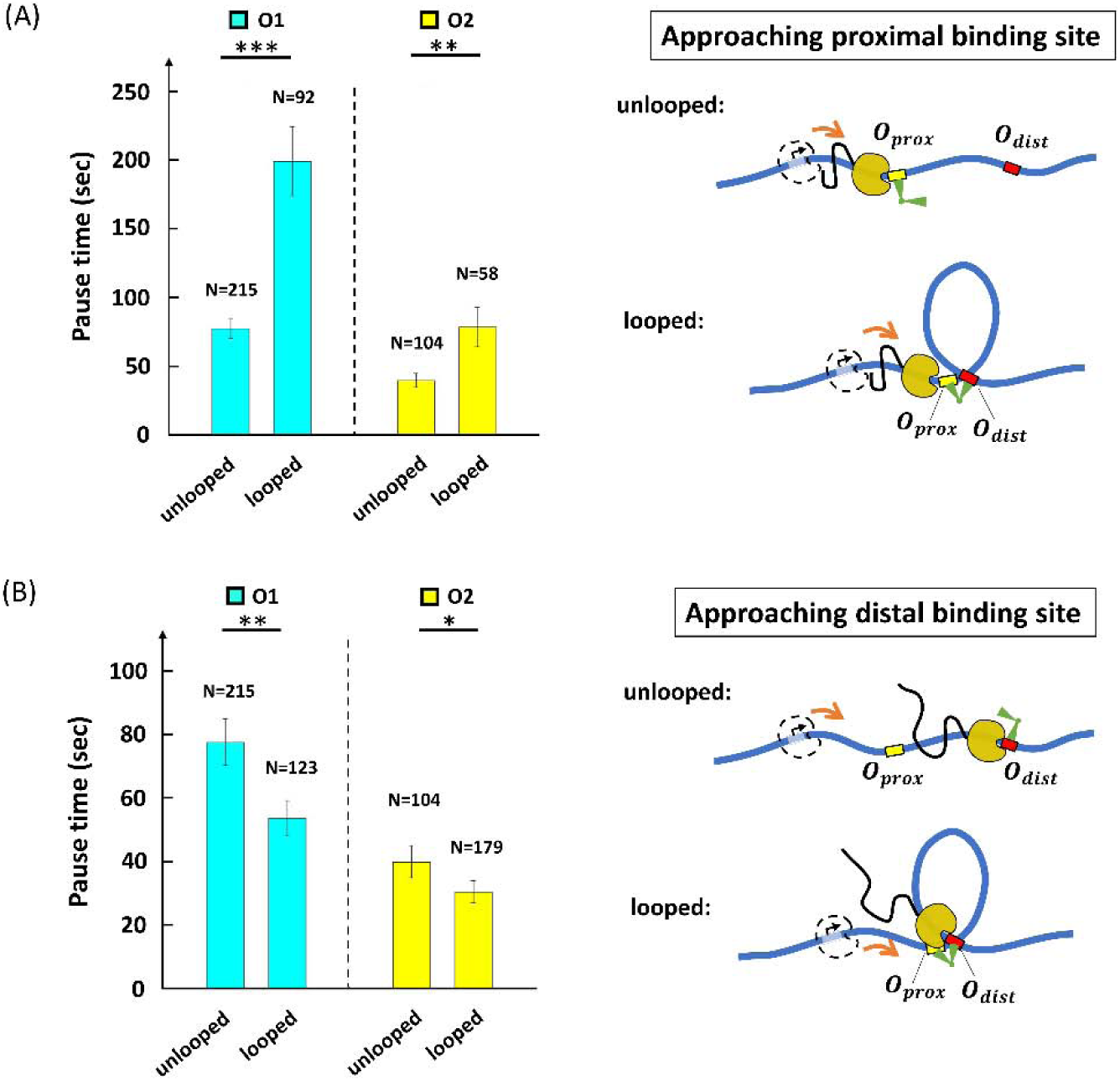
The LacI-mediated loop enhances and attenuates RNAP pausing at the proximal and distal binding sites respectively. (A) Average pause durations were longer at the proximal operator consisting of either O1 (cyan) or O2 (yellow), in the looped with respect to the unlooped conformations depicted at right (\*\*\****p*** ≤ **0. 001**, \*\****p*** ≤ **0. 01** for two-sample t-tests). (B) Average pause durations were shorter at the distal operator consisting of either O1 (cyan) or O2 (yellow), in the looped with respect to the unlooped conformations depicted at right (\*\****p*** ≤ **0. 01**, * ***p*** ≤ **0. 12** for two-sample t-tests). Standard errors and numbers of samples, *N*, are indicated.

Pauses at LacI-O1_prox_ obstacles were 77 ± 7 s (N=215) without a loop, but 199 ± 25 s (N=92) with a loop. Pauses at LacI-O2_prox_ obstacles were 40 ± 5 s (N=104) without a loop, but 79 ± 14 s (N=58) with a loop. A likely reason for the increases is that the secondary, distal binding site increases the local concentration of LacI and the effective affinity for the proximal binding site (27). The steric hindrance caused by the loop itself may also contribute to the lengthening of the pause at the proximal binding site, although this effect is likely to be small (33). It is also informative that looped LacI-O1_prox_ obstacles obstructed transcription for a considerably longer time (199 s / 77 s) than looped LacI-O2_prox_ obstacles (79 s / 40 s) with respect to unlooped templates. Since LacI has higher affinity for the O1 than the O2 binding site, LacI is more likely to remain at O1 than at O2 after the loop break down, increasing the probability that an elongating RNAP may pause at O1.

We next investigated RNAP approaching the promoter-distal operator, O_dist_. When RNAP is at the distal operator, the tether length will not be changed significantly by loop formation, because it does not change the segment ahead of RNAP (Figure S1A and cartoon (V) on the right of Figure 1C). We used traces such as the ones in the rightmost panel of Figure 1B and in interval (V) of Figure 1C to compare pause durations at O_dist_ (Figure 1C, interval V; Figure S6, interval V), with or without looping. Under the 0.2 nM LacI concentration used, 41% of the LacI obstacles at O_dist_ can be assumed to secure loops (Figure S5). Surprisingly, and in contrast to O_prox_, we observed that under these conditions, the average pause time of RNAP at LacI-O1_dist_ and LacI-O2_dist_ was shortened with respect to obstacles on unlooped templates to 54 ± 6 s (N=123) and 30 ± 4 s (N=179), respectively. Assuming 41% looped obstacles, RNAP inside the loop pauses at LacI for approximately 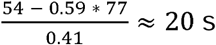 s at the distal O1 operator, and for 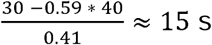 s at the distal O2 operator. These results suggest that transcription within the loop promotes the release of LacI from a distal operator site, despite the locally increased concentration of LacI and regardless of the operator affinity.

### Transcription of looped segments is slower

Inspection of interval IV in Figure 1C suggests that a protein-mediated loop can significantly delay transcription by RNAP. According to the twin-supercoiled-domain model (28), rotation of the DNA template unwinds DNA behind the transcribing RNAP, generating negative supercoiling (Figure 3A, red DNA), and winds DNA ahead, generating positive supercoiling (Figure 3A, yellow DNA). Furthermore, since a LacI-mediated loop constitutes a topological domain (45), transcription within the loop will generate torsional stress. Within a 400 bp-long loop, the torsional stress created by a transcribing RNAP can quickly accumulate to +11 pN·nm ahead or -11 pN·nm behind, stalling RNAP progress(42). We estimate that RNAP might translocate as few as four bp within the 400 bp loop before stalling (Materials and Methods). Stalled RNAP is prone to backtracking, and recovery from the backtracked state delays RNAP to slow transcription. Thus, we measured the total time required to transcribe the loop segment in each trace (duration of interval IV in Figure 1C or intervals such as IV in Figure S6), averaged it over all traces in the presence of LacI/looping and compared it with the average time required to transcribe between the two operators in the absence of LacI/looping (Figure 1B, left). The average transcription time for looped O1-O2 and O2-O1 segments were 192 ± 31 s (N=104) and 185 ± 29 s (N=86), respectively. Both these times were much longer than the average time without looping (32 ± 5 s; N=35). We conclude that the LacI loop can significantly hinder RNAP progress by generating torsional stress.

**Fig. 3.**
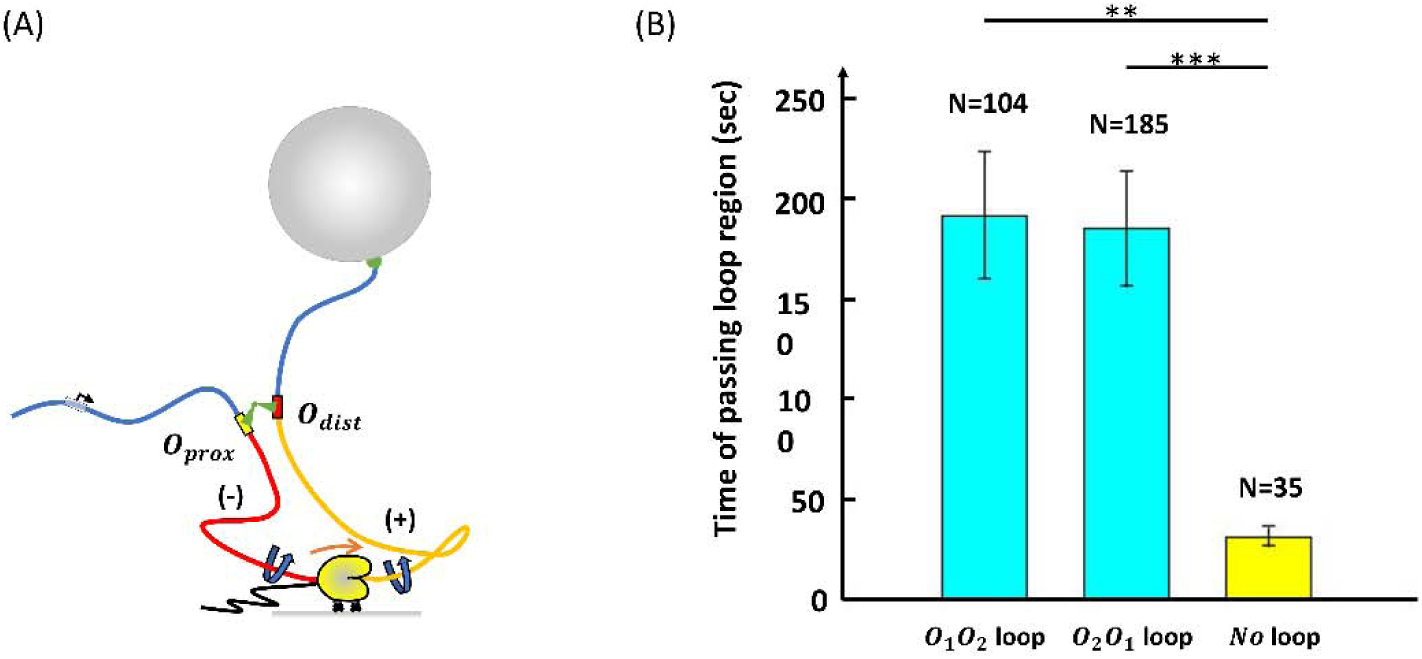
RNAP transcribes a loop more slowly. (A) A cartoon depicting RNAP transcribing a loop. The right-angle black arrow indicates the promoter. The blue-colored DNA segments are torsionally relaxed. The red-colored DNA segment is unwound, while the yellow DNA segment is overwound by RNAP. Nascent RNA is the thin, black line emerging from RNAP. (B) RNAP requires almost tenfold more time to transcribe a looped segment (cyan) compared to the same unlooped segment (yellow). \*\*\****p*** ≤ **0. 001**, \*\****p*** ≤ **0. 01** (two-sample t-tests). Standard errors and numbers of samples, *N*, are indicated.

### RNAP surpasses LacI obstacles faster on positively supercoiled templates

The strength of the LacI roadblock may be affected not only by loop formation, but possibly also by accumulated torsional stress within the loop. In particular, the data in Figure 2 showed that the loop alleviates the RNAP obstruction by the distal LacI obstacle. We hypothesized that the shorter pauses at the distal operator might result from destabilization of LacI-O_dist_ complexes by transcription-generated positive supercoiling. In the TPM experiment, the DNA segments flanking the loop (blue in Figure 3A) quickly swivel about single covalent bonds to the attached surfaces to release any torsional stress due transcription (Figure S1A). Within the loop, however, positive supercoiling ahead of RNAP accumulates and might cause LacI to dissociate from the DNA. To test this hypothesis, we used magnetic tweezers (MTs) to follow elongation of an RNAP transcribing toward a LacI-O1 obstacle on a construct where the DNA ahead could be positively supercoiled (Figure 4A, Figure S1B). In this experiment, the segment between RNAP and the tethered bead was rotationally immobilized by multiple biotin-streptavidin linkages to the bead. The promoter of this template was 252 base pairs (approximately 24 turns) upstream from the O1 binding site (Figure S1B). To create positive supercoiling just as RNAP arrived at the LacI obstacle, the DNA template was preloaded with negative plectonemes (Figure 4A) under forces ranging between approximately 0.25 and 0.8 pN. As RNAP transcribes, the DNA tether length would be expected to change as depicted in Figure 4A. We first verified that RNAP could transcribe a tether preloaded with -24 turns (gray and black trace in Figure 4B). After introducing NTPs (blue arrow in Figure 4B), the tether extension increased, due to annihilation of the negative supercoiling by the positive supercoiling generated by the elongating RNAP, until the DNA tether became torsionally relaxed. After that, RNAP continued to wind the DNA tether introducing positive plectonemes until the bead was drawn to the surface of the flow-chamber, or RNAP stalled due to either steric hindrance by the plectonemes, or perhaps by large torsional stress. Once the ability of RNAP to transcribe a negatively supercoiled template by several turns was verified (Figure 4B, grey and black curves), transcription was recorded in the presence of LacI after pre-loading the template with -19 turns. RNAP was expected to reach the O1 binding site after having supercoiled the DNA template to ΔLk = +5, according to extension versus twist curves (Figure S4) in which the tether length with +5 turns can be clearly distinguished from that of the same torsionally-relaxed tether (ΔLk = 0). The blue and red traces in Figure 4B are the raw data and a 4 s moving average, respectively, of a measurement with 10 nm LacI. Although RNAP paused also at random positions along the trace, the expected pause at O1 was clearly distinguishable (Figure 4B, between vertical, black, dashed lines, see Figure S4 for identification of the O1 position). For comparison, we acquired MT measurements of the RNAP pause in front of the LacI-O1 obstacle on torsionally relaxed (nicked) DNA under 0.45 pN tension (Figure 4C right, Figure S1B).

**Fig. 4.**
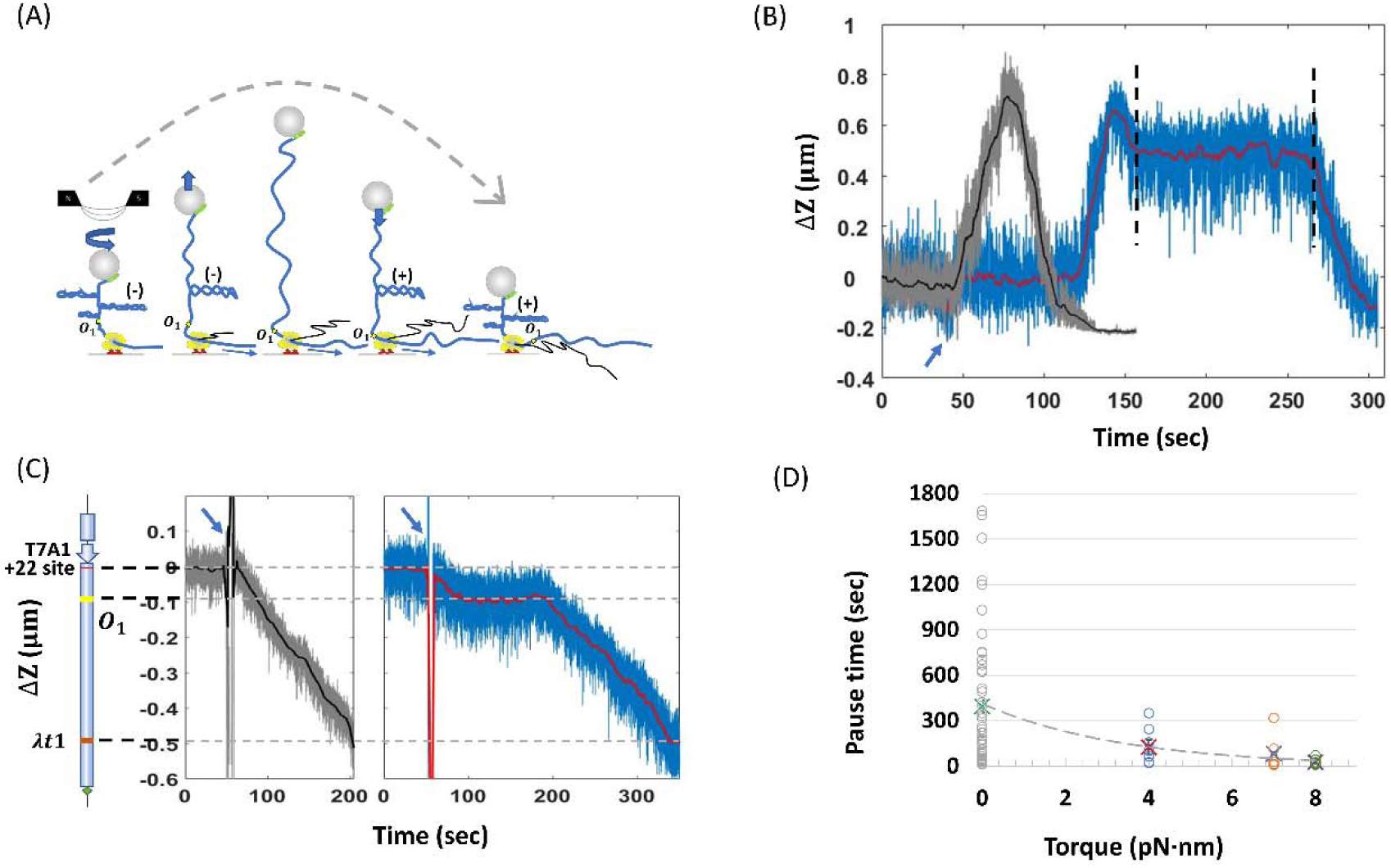
Comparison of RNAP pause times at O1 with and without positive supercoiling. (A) A DNA tether was mechanically unwound forming plectonemes prior to the addition of NTPs. Subsequent transcription introduced positive supercoils that annihilated the mechanically induced, negative supercoils and lengthened the tether to a maximum. Further transcription and positive supercoiling eventually produced plectonemes that contracted the tether length. The dashed gray curve indicates the extension of the DNA tether during progressive conversion of negative to positive plectonemes due to transcription by RNAP. (B) Representative observations of extension versus time were recorded during transcription without (gray) and with (blue and red) LacI on a template pre-loaded with negative supercoiling. The blue arrow indicates the time at which all four NTPs or NTPs+LacI were introduced. The two vertical black dashed lines circumscribe a pause by RNAP at the LacI-O1 operator complex. (C) Representative observations of elongation were recorded in the absence of pre-loaded supercoiling without (left panel, gray and black) or with (right panel, blue and red) LacI. The cartoon on the left of the two traces is a schematic representation of the DNA template (D) Pause times by RNAP at the LacI obstacle varied as a function of torque on the DNA. Torque values were calculated using Eq 1 in reference 38, except that C_eff_ was estimated in 100 mM [Na+] instead of 50 mM [K+]. Circles represent measured pause times, while crosses represent the average pause times.

Figure 4D shows how pause times changed with different torque on the DNA. The average RNAP pause at the LacI-O1 obstacle with no torque (τ _1_ = 0 pN·nm) (data: gray circles, average: green cross) was 393 ± 64 s (N=49), much longer than the average pause time with positive torques τ _2_ ∼ 4 pN·nm (data: blue circles, average: red cross), τ _3_ ∼ 7 pN·nm (data: orange circles, average blue cross) and τ _4_ ∼ 8 pN·nm (data: green circles, average: purple cross), which pause RNAP for 125 ± 42 s (N=8), 82 ± 50 s (N=6) and 26 ± 8 s (N=8), respectively. Thus, positive supercoiling significantly facilitated transcription through the Lac-O1 obstacle and likely disrupted the LacI-O2 obstacle as well. The decrease in pause duration as positive torque on the DNA increased suggests that positive supercoiling weakens LacI binding and transcriptional roadblocking. Positive supercoiling generated by RNAP translocation is also known to destabilize nucleosomes (32, 46); it may, therefore, represent one means by which RNAP removes protein roadblocks along the DNA.

## Discussion

Here, we present evidence that the ability of DNA-bound LacI to act as barrier for the transcribing RNAP is strongly dependent on DNA topology. Our TPM measurements show that a DNA loop formed by LacI bridging two operator sequences alters the roadblocking effect of LacI-operator obstacles in opposite ways depending on their position relative to the promoter (proximal vs. distal). Approaching a loop from the outside, RNAP paused in front of LacI-proximal operator roadblock longer than it did when there was no loop. Such increases in pause duration are likely due to the fact that the loop effectively increases the LacI concentration in the vicinity of the operator (27). This increases the occupancy of the O_prox_ binding site and may sterically hinder approaching RNAP (33). Once RNAP clears the proximal operator, it is dramatically decelerated within the 400 bp looped segment, as compared to transcription of the same DNA in an unlooped configuration, likely as a result of torsionally induced stalling. Yet, once RNAP transcribes toward the end of the loop, it clears the distal LacI roadblock faster than the same roadblock in the absence of DNA loop.

Torsional disruption of LacI-DNA complexes might explain the shorter pauses at distal LacI securing a loop. Indeed, using magnetic tweezers to arrange positive supercoiling of the DNA to coincide with the RNAP arrival at the distal LacI obstacle shortened pauses by RNAP. This is strong evidence that positive supercoiling generated by transcription facilitates clearance of the LacI obstacle. In general, accumulated positive supercoiling ahead of RNAP may accelerate protein dissociation from DNA and shorten pauses at protein-mediated loops or other DNA structures.

Further control experiments using topoisomerase IB, or a nicking enzyme targeted to the loop region to artificially release the accumulating torsion, were not productive due to the inability to synchronize activities of those enzymes with RNAP elongation and nicking of either DNA strand induced RNAP pausing, or undesired transcription initiation, at nicks. Nonetheless, the results reported in this work strongly indicate that small loops of few hundred base pairs, such as the one considered here and those induced by many prokaryotic regulators, significantly slow transcription by RNAP, and that the positive torsional stress accumulated ahead of a transcription elongation complex may help to clear the path.

Generally, destabilization of DNA roadblocks as positive supercoiling accumulates is likely to enhance RNAP progression along DNA with bound proteins. Simultaneously, negative supercoiling trailing the transcription complex may help dislodged proteins rebind and/or may stabilize proteins behind the complex. *In vivo*, transcription would generate supercoiling at rates of 3.9 - 5.5 turns/sec (39 – 55 bp/s) (47), a potent source of supercoiling for topoisomerases to manage. Looping transcription factors that can shift between sites ahead and behind transcription complexes would maintain at least one connection to the DNA as RNAP passes and avoid diffusing away from their binding sites. This adds a fine level of control to that exerted by the overall concentration of the transcription factors.

*In vivo*, the ability of protein-mediated loops to hinder RNAP elongation may be a critical factor in the regulation of transcription at the local level. In the eukaryotic organism *Drosophila melanogaster*, it was observed, using 4C-seq assays, that RNA polymerase II was often paused near promoters involved in long-range interactions via several kilobase pair-long loops with enhancers. The authors hypothesized that since promoter-proximal complexes can exert enhancer-blocking activity (48), the presence of paused polymerase could safeguard against premature transcriptional activation, and yet keep the system poised for activation (49). It is possible that in the case of regulatory loops found along the template during elongation, RNAP pausing, either in front or within, is part of a mechanism to (i) wait for additional factors to resolve the loop, or relieve supercoiling, as a signal to restart transcription, or (ii) avoid over transcription of a gene. Given the ubiquity of looping in any genome, we propose that RNAP may temporarily stall inside loops where it may be able to respond to regulatory factors and eventually transit the loop segment dispersing protein-DNA obstacles through the generation of positive supercoiling. In this way, RNAP can carve its path through DNA structures and DNA-bound proteins that are necessary for genome maintenance and regulation of gene expression in all life.

## Supplementary Figures

Please, see accompanying SI file

## Acknowledgments

LacI was a generous gift from Kathleen Matthews, Rice University. We thank Karen Adelman, Harvard, for generously providing RNAP for initial experiments.

This work was supported by the National Institutes of Health (NIH) grants R01 GM084070 to LF and R01 GM067153 to IA.

## Author contributions

W.X. Contributed measurements, to the design of experiments, and to writing of the paper

Y.Y. Contributed measurements

I.A. Contributed RNA polymerase and to writing of the paper

D.D. Contributed to the design of experiments and to writing of the paper

L.F. Contributed to the design of experiments and to writing of the paper

## Data Availability

Data will be made available on Dataverse at Emory.

