## Supplementary material for "Positive supercoiling favors transcription elongation through lac repressor-mediated DNA loops": Figures S1-6 and TableS1

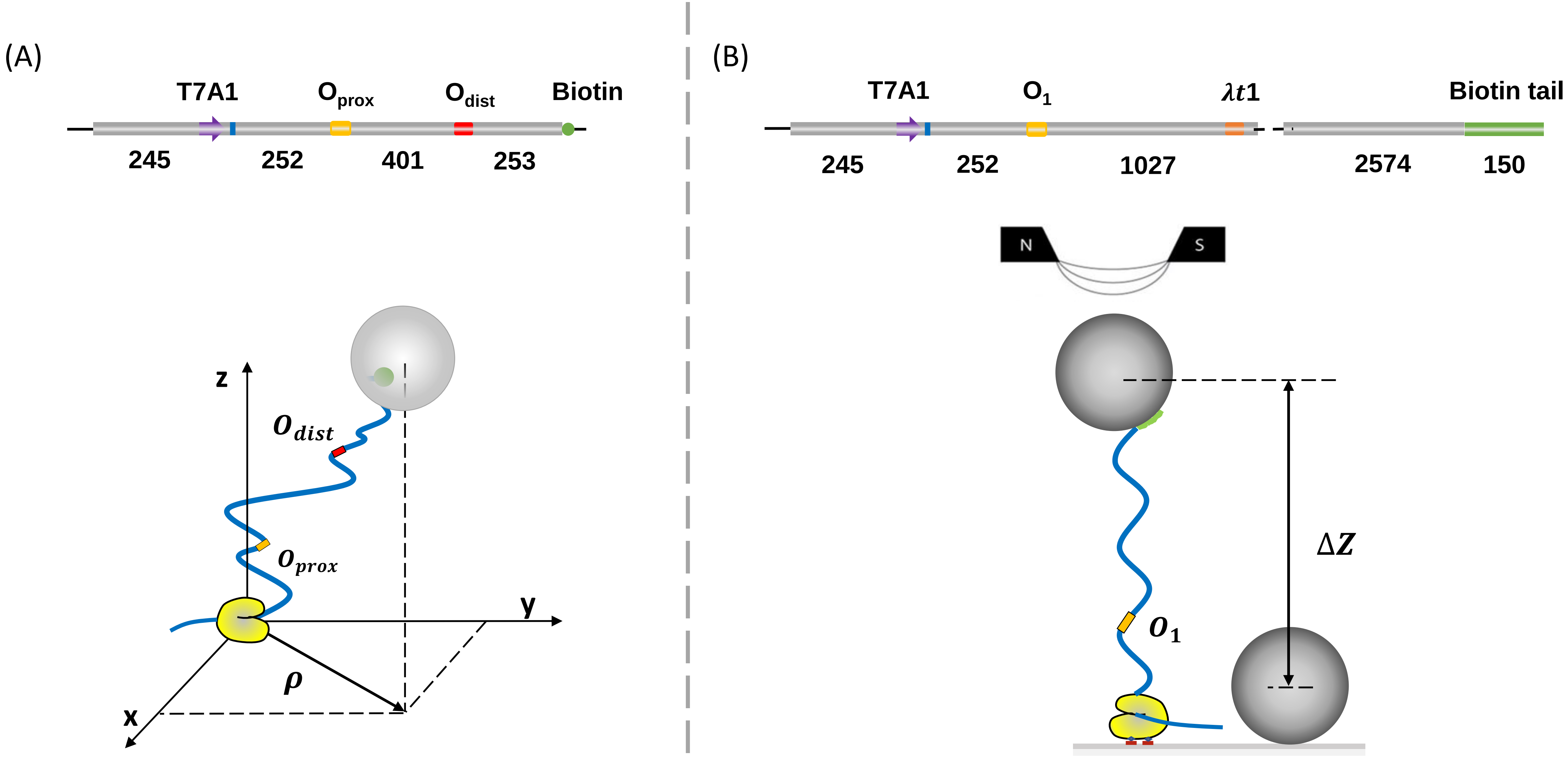

(A)
T7A1
O prox
O dist
Biotin
245
252
401
253
z
O dist
O prox
y
ρ
x
(B)
T7A1
O 1
λt1
Biotin tail
245
252
1027
2574
150
N
S
ΔZ
O 1

**Fig. S1. Experimental construct and setup for TPM/MT measurements.** RNAP anchored the DNA template to the glass surface of a microchamber. Both DNA templates contained a T7A1 promotor for RNAP binding and a stall site at position +22. (A) A streptavidin-coated polystyrene bead labelled the biotin-tagged free end of DNA molecules containing two binding sites for the lac repressor. The variation in time of the 2-D projection of the bead served as a reporter of the Brownian motion of the bead. (B) A streptavidin-coated super paramagnetic bead bound to a biotin tail tagged the free end of DNA molecules containing the O1 binding site for the lac repressor. The motion of a tethered bead relative to that of a stuck bead served to assess the changes in height of the bead.

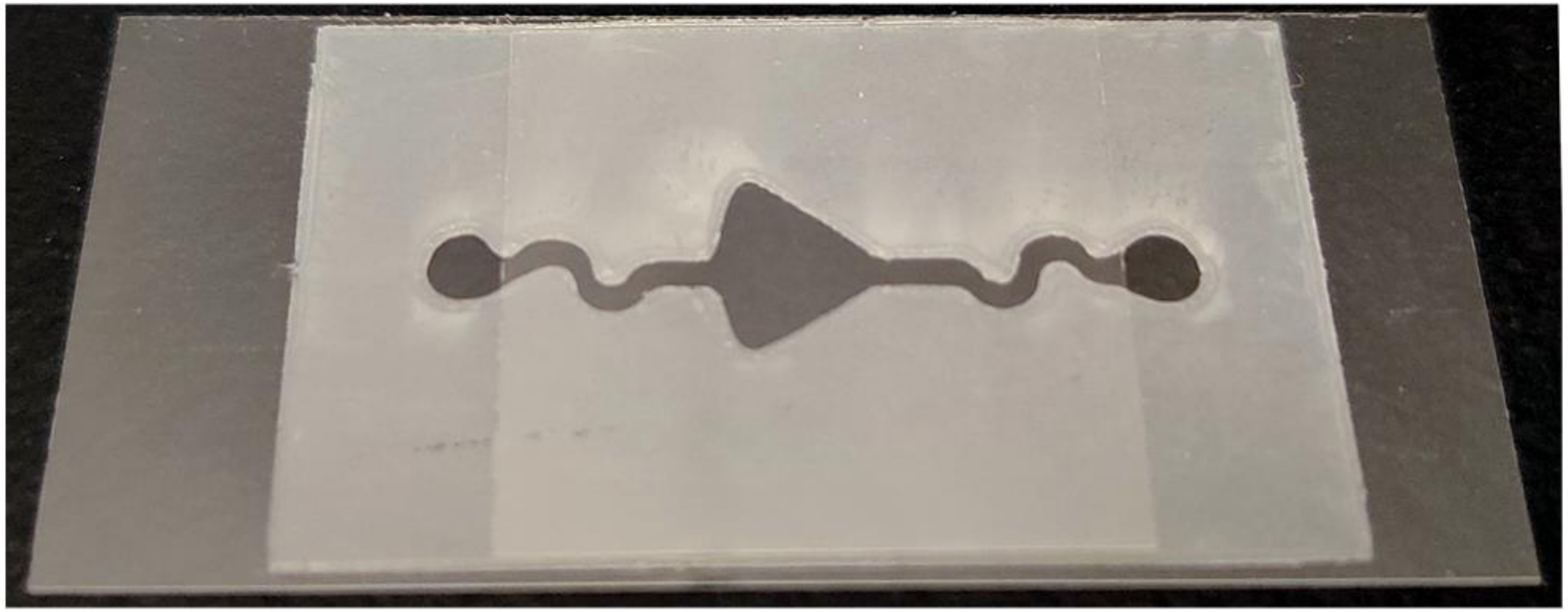

**Fig. S2. Image of a typical micro chamber.** The bottom coverslip supports a parafilm gasket with inlet and outlet reservoirs at the edges of the top coverslip connected through narrow inlet and outlet channels to the central observation area under the coverslip. The parafilm and coverslips are heated and sealed together, and the total volume of the chamber is approximate 6 $\mu$L.

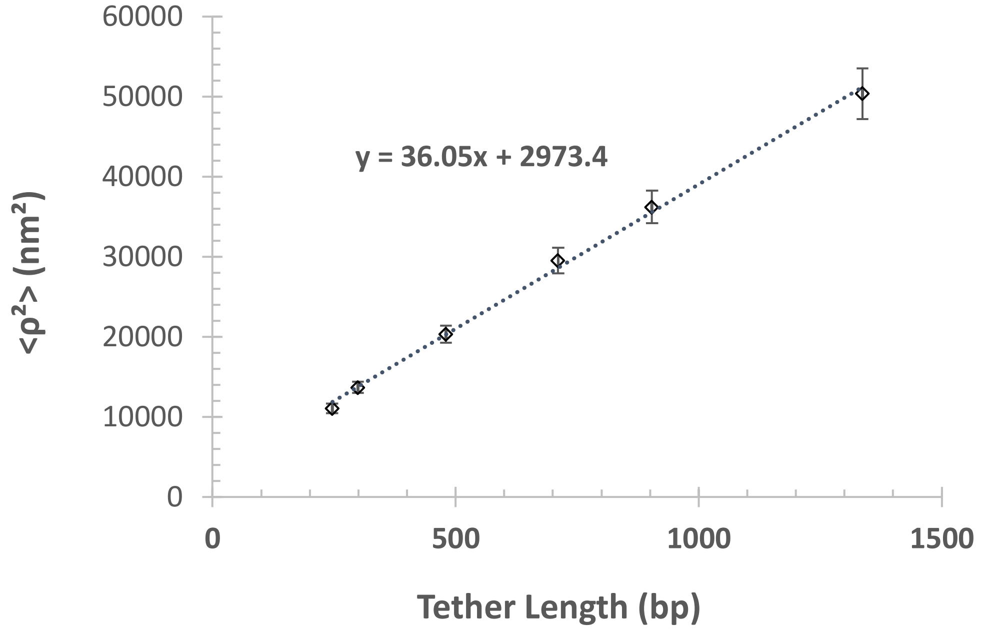


| Tether length (bp) | $<\rho^2>$ $(nm^2)$ |
|---|---|
| 246 | 11075 |
| 299 | 13680 |
| 480 | 20345 |
| 711 | 29508.5 |
| 904 | 36228.3 |
| 1337 | 50375 |

**Fig. S3. Calibration of tether length and rho square.** The recorded rho square values were converted into values of tether length by measuring rho square for tethers of different known lengths (244, 299, 480, 711, 904, 992, 1337 bp). The expression for the fitted line is: $< \rho^2 >= 36.05 * \text{tether length} + 2973.4$.

**Fig. S4.**

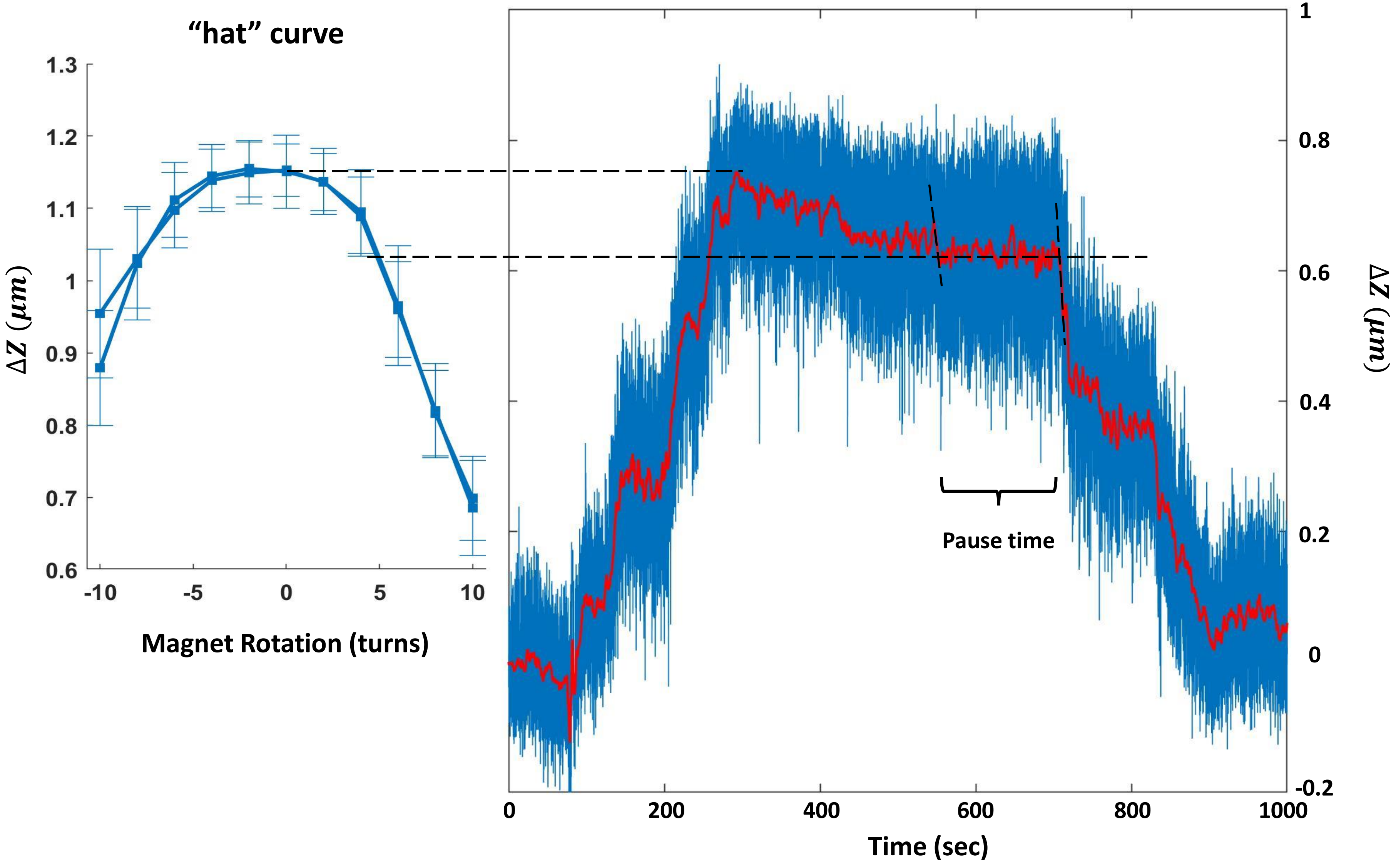

“hat” curve
1.3
1.2
1.1
1
0.9
0.8
0.7
0.6
$\Delta Z\ (\mu m)$
-10
-5
0
5
10
Magnet Rotation (turns)
1
0.8
0.6
0.4
0.2
0
-0.2
$\Delta Z\ (\mu m)$
0
200
400
600
800
1000
Time (sec)
Pause time

**Fig. S4. Measurement of pause time on supercoiled DNA.** The curve on the left is the "hat" curve which represents the relationship between tether extension and magnet rotation. The trace on the right is one representative trace that starts transcription after -19 turns had been introduced in the DNA template of a stalled TEC. The two horizontal, black dashed lines show that the elongation pause occurred after RNAP had introduced +5 turns in the template. Given the distance of 253 bp between the stall and the O1 site, this is expected to be exactly in front of O1: [(253bp/10.4bp/turn) – 19 turns] = ~5 turns.

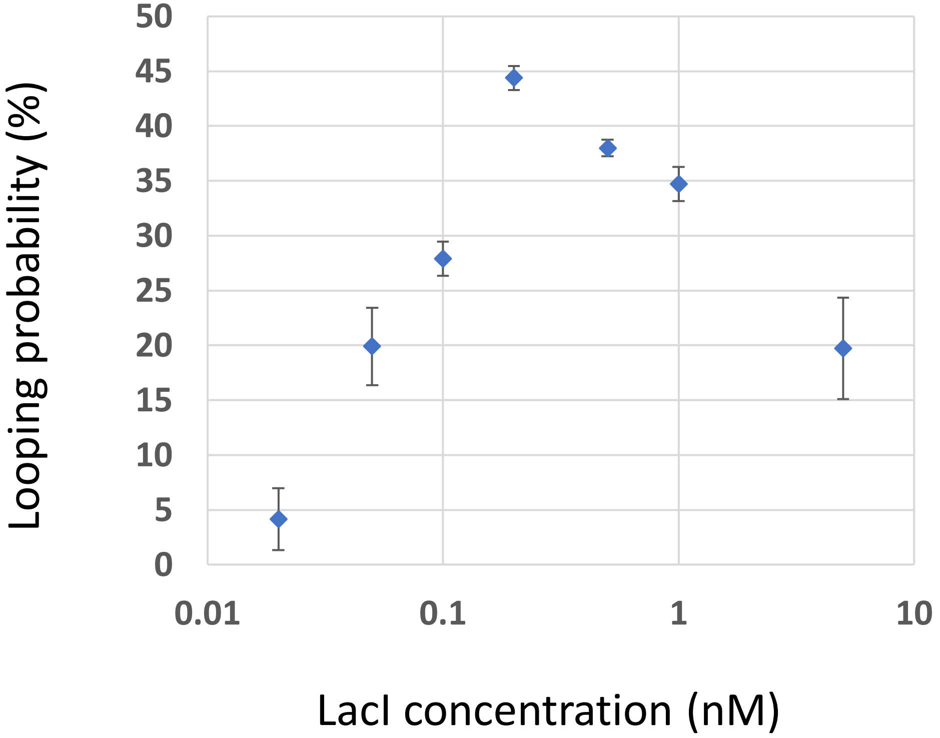


| [LacI] (nM) | Looping Probability | Stand_Error |
|---|---|---|
| 0.02 | 0.041461 | 0.028293 |
| 0.05 | 0.199028 | 0.035377 |
| 0.1 | 0.278917 | 0.015676 |
| 0.2 | 0.443856 | 0.010945 |
| 0.5 | 0.379909 | 0.007482 |
| 1 | 0.3471 | 0.015456 |
| 5 | 0.197158 | 0.046178 |

**Fig. S5. Looping probability vs LacI concentration.** The looping probability was measured in 904 bp-long DNA tethers with a 400 bp loop region under 0.02, 0.05, 0.1, 0.2, 0.5, 1 and 5nM LacI concentration in transcription buffer (TXB).

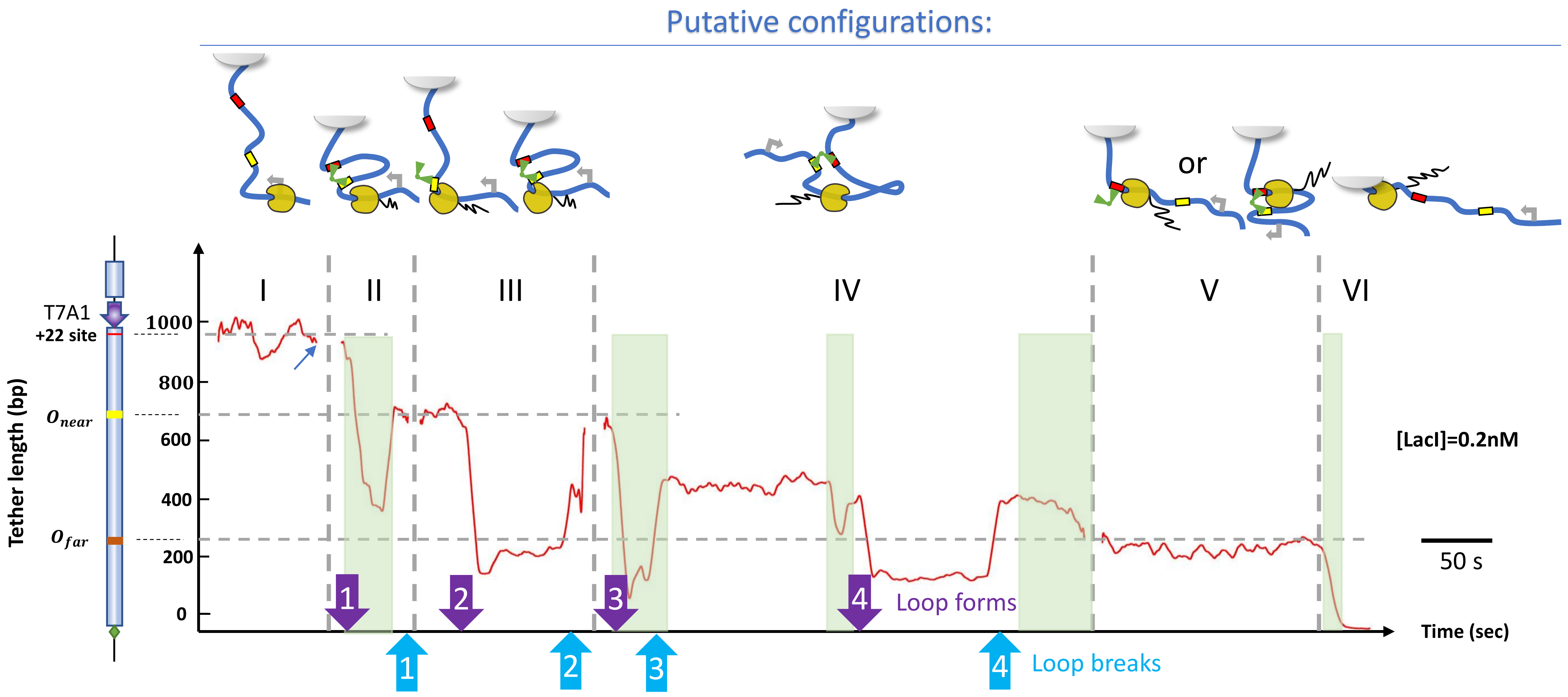

Putative configurations:
or
I
II
III
IV
V
VI
T7A1
+22 site
$O_{near}$
$O_{far}$
Tether length (bp)
1000
800
600
400
200
0
[LacI]=0.2nM
50 s
Time (sec)
1
2
3
4
Loop forms
1
2
3
4
Loop breaks

**Fig. S6.** A LacI-mediated loop can produce discrete jumps or plateaus as transcription decreases the tether length. To facilitate interpretation, an illustrative example was assembled from segments drawn from different traces. Note that transcription draws the bead toward the anchor point and shortens the tether. Intervals (I) (II) (V) (VI) are similar to the intervals (I) (II) (V) (VI) in the representative trace of Fig. 1C. The green areas indicate transcriptional activities, the purple arrows (PA) mark the time that loop forms while the blue arrows (BA) mark the time that loop breaks. In interval (I), the DNA tether length remains constant before introducing NTPs and LacI (red arrow). Transcription begins shortly after introducing NTPs in interval II and a loop forms (PA 1), but the loop ruptures (BA 1) as RNAP approaches Onear and RNAP pauses at this location suggesting that LacI is bound to Onear. In interval (III), a loop forms (PA 2) while RNAP is paused in front of $O_{near}$ and ruptures (BA 2) shortly thereafter, suddenly restoring the tether length to $O_{near}$. At the beginning of interval (IV), RNAP begins to transcribe the inter-operator region downstream of $O_{near}$ as a loop forms (PA 3). Loop breaks down (BA 3) after about 40 seconds and the new tether length corresponds to RNAP approximately in the middle of the loop segment, 200 bp from $O_{near}$. Then, RNAP stalls for 150 second until the loop reforms (PA 4). Judging from the tether length at the time loop (BA 3) ruptures, RNAP has not progressed further while circumscribed by loop. Transcription resumes shortly after loop ruptures (BA 4) and then RNAP pauses at $O_{far}$, interval (V). This indicates that LacI is bound to $O_{far}$. In this region, loop formation may occur but does not significantly change the tether length and cannot be detected. In interval (VI) RNAP has passed $O_{far}$ and transcribes to the end of the DNA template.

Table S1. **plasmids and primers.**

| Construct | O1-400-O2 (TPM) | O2-400-O1 (TPM) | O1 (MT) | Bio-, dig-tails |
|---|---|---|---|---|
| Template | pWX_12_400 | pZV_21_400 | pRS_1N_400 | pBluKSP |
| Forward primer | S/JBOIDO1_400/2086 | S/JBOIDO1_400/2086 | S/JBOIDO1_400/2086 | A/pIT_Loop3/1665 |
| Reverse primer | B-A/YY400/3342 | B-A/YY400/3342 | A/pO1O2_401/3043_ApaI | Pol1979R |
| Restriction enzymes | | | ApaI | ApaI |

| Prime name | Sequences |
|---|---|
| S/JBOIDO1_400/2086 | agcttgtctgtaagcggatg |
| B-A/YY400/3342 | [bio]-accgcatcagcaagtgtat |
| A/pO1O2_401/3043_ApaI | ctgggcccggtgaatccgttagcga |
| A/pIT_Loop3/1665 | ggcgattaagttgggtaacg |
| Pol1979R | tgtggaattgtgagcggata |

**Table S1. Plasmids and Primers.** Plasmids and primers used to produce DNA segments in this work (Top). Primer sequences. Bio- means biotinylated at 5’ end of primer (Bottom) .
